# Unique deficits in place coding across subfields of the hippocampus in a mouse model of temporal lobe epilepsy

**DOI:** 10.64898/2025.12.17.694941

**Authors:** Brittney L. Boublil, Margaret M. Donahue, Cathy B. Dang, Gergely Tarcsay, Laura A. Ewell

**Affiliations:** Anatomy and Neurobiology, School of Medicine, University of California, Irvine, Irvine, CA, USA; Neurobiology of Brain and Behavior, Charlie Dunlop School of Biological Sciences, University of California, Irvine, Irvine, CA, USA; Center for Learning and Memory, University of California, Irvine, Irvine, CA, USA

## Abstract

**Objective:** Memory problems are comorbid with Temporal Lobe Epilepsy (TLE). Animal models of TLE reveal impairments in spatial firing fields of hippocampal place cells, providing a potential neural substrate for memory problems. Each subfield of the hippocampus carries out unique aspects of spatial memory, yet little is known about how individual subfields are perturbed. Here, we investigated the spatial coding properties of the three major subfields of the hippocampus.

**Methods:** Single unit recordings were made from CA1, CA3 and the dentate gyrus (DG) of mice (N = 10, 6M/4F) induced with epilepsy using the supra-hippocampal kainate model and in control mice injected with saline (N = 6, 3M/3F). Place cell activity was measured while mice foraged in highly familiar environments to assess basic place cell properties and in novel environments to assess remapping.

**Results:** A lower percentage of cells were classified as place cells in CA1 of epileptic mice, whereas percentages were similar in CA3 and DG compared to control. Place fields of CA1 were less coherent, place fields of CA3 were less stable, and place fields in DG has smaller differences between in-field and out of field firing. All regions constructed new distinct maps within the first session of exposure to a novel environment, however new maps in CA3 trended toward instability.

**Significance:** These results point to specific deficits within subfields of the hippocampus, which may indicate that there are different cellular and network mechanisms at play. Such heterogeneity would be predicted to contribute differently to memory deficits.

**Key Points:**

- CA1 exhibits place map quality problems; CA1 has fewer place cells and those remaining have reduced spatial coherence
- CA3 exhibits place map stability problems; CA3 has lower spatial correlation within sessions and trends toward not forming stable new maps in a novel environment.
- DG exhibits reduced signal to noise; DG has smaller differences between in-field and out of field firing rates.

## 1 INTRODUCTION

Memory comorbidity is common in epilepsy and greatly impacts patient’s quality of life (1, 2). Disease associated injury to temporal lobe structures is commonly thought to be the culprit. Recent work, however, highlights the role of altered network function during interictal periods (3), which is significant because network function is a rescuable target. In the case of temporal lobe epilepsy (TLE), it is well established that behaviorally relevant firing patterns of hippocampal place cells (neurons with spatially tuned firing patterns) are disrupted in multiple rodent models (4, 5, 6, 7, 8, 9). Disrupted spatial coding of hippocampal place cells is a strong candidate for the neural substrate of impaired memory in TLE.

The hippocampus comprises several subfields – with each providing distinct roles in spatial memory (10, 11). Signals enter the hippocampus at the dentate gyrus (DG) and then pass on to area CA3 and finally to area CA1, which sends outputs to subiculum and cortex. Each region processes the signal – associating components of the memory together into one representation and putting it into a form suitable for long-term storage in the cortex. The DG is a sparsely activated structure thought to perform pattern separation (12, 13) and novelty detection (14). Thus, the DG transforms the cortical signals into maximally distinct representations and ‘teaches’ the downstream CA3, potentially highlighting when representations are novel and thus tagged to be learned. Area CA3, which sits at the core of hippocampal processing, has unique anatomy compared to the other hippocampal subfields – it is a highly recurrent structure (i.e., auto-associative) in which neurons receive input from their own axons, and commissural connections from the contralateral CA3 (15). It is proposed that such recurrence supports the storage and recall of ensemble representations of distinct features of environments (separated by the DG) (16) and supports temporal ordering of experience (17). CA1 receives the signal from CA3 and adds additional salient features to the population representation such as environment-associated reward outcomes (18) or goal locations(19). CA1 indexes the hippocampal memory back out to cortex during slow wave sleep (20, 21).

In TLE, all sub-regions of the hippocampus exhibit pathology. The DG undergoes massive rewiring and experiences significant inhibitory interneuron loss (22, 23, 24, 25, 26, 27). The CA fields experience sclerosis (28). Despite these widespread changes across the hippocampal subfields, the overwhelming majority of studies of place coding in epilepsy have focused on area CA1 exclusively (4, 5, 6, 7, 8, 9), and have been carried out in systemic models of temporal lobe epilepsy that likely impact regions beyond the hippocampus. We performed single unit recordings in freely moving mice, targeting the three major subfields of the hippocampus. Employing the supra-hippocampal kainic acid model of chronic TLE, we tested how TLE impacts place coding across subfields. We have previously reported that mice with epilepsy induced with this model experience frequent subclinical seizures and interictal spiking as well as deficits in object location memory and spatial working memory (29) (30). Here, we recorded while mice foraged in highly familiar environments and during exploration of a novel environment. Familiar environment recordings provided a baseline understanding of region-specific place codes, while novel room recordings allowed us to assess the development and stability of new maps.

## 2 MATERIALS AND METHODS

### 2.1 Mice

A total of 16 C57BL/6 mice (nine males and seven females) were used for this study. Mice were individually housed on a reversed 12/12-hour light/dark cycle. The weights of all mice were maintained on an *ad libitum* diet except during the open field foraging task, when mice were food restricted to maintain 85% of their baseline weight. All experimental procedures were approved by the Institutional Animal Care and Use Committee (IACUC) at the University of California, Irvine.

### 2.2 Induction procedure for supra-hippocampal kainate model

All surgeries were performed using aseptic procedures. Mice (12 weeks old) were initially anesthetized with isoflurane gas and transferred to a Kopf stereotaxic instrument, where they were positioned and secured with an anesthesia mask and non-rupture ear bars. Anesthesia was maintained throughout the surgery at 1-2% isoflurane delivered in O_2_ at 1 L/min and body temperature was maintained at 37–38°C using reusable gel heat packs. Eyes were protected with a lubricating eye ointment. Carprofen (Patterson Veterinary, 5 mg/kg) was administered subcutaneously as an analgesic. The scalp was prepared by removing the hair and washing the scalp with iodine. Lidocaine (Patterson Veterinary, 4 mg/kg) was injected subcutaneously under the scalp, and a midline incision was made. The skin was retracted, and a craniotomy was made above the hippocampus (AP: −2.0 mm, ML: +1.5 mm from bregma) using a carbide bur.

Kainic acid (KA) (Tocris) was dissolved in saline to provide a solution with a concentration of 20 mM (31, 32, 33, 34). A glass pipette was lowered into the brain (DV: −0.8 mm from dura) and left in place for a minimum of 1 minute before the injection. 70 nL of KA (or sterile saline for controls) was injected using a custom microinjection system (Narishige International, #MO-10), over the span of 2 min. The glass pipette remained in place for an additional 1 minute following the injection to prevent the backflow of the solution up the needle tract. The incision was closed with absorbable sutures. *Status epilepticus* was terminated 4 hours later with a subcutaneous injection of either lorazepam (West-Ward, 7.5 mg/kg) or diazepam (Patterson Veterinary, 20 mg/kg). Lorazepam/Diazepam was not injected for saline control mice.

### 2.3 Implantation of Open Ephys ShuttleDrive

Four weeks after *status epilepticus*, mice were implanted with an Open Ephys ShuttleDrive (35), which is composed of 16 individually moving tetrodes. Surgical preparation was the same as described above. Once anesthetized and secured to the stereotaxic instrument, buprenorphine (Patterson Veterinary, 0.1 mg/kg) was administered subcutaneously as an analgesic. Once the skull was exposed, additional surface area was created to improve the bonding of the dental composite by scoring the skull bone and by making shallow divots on the skull’s surface using a carbide bur. Optibond was applied over the prepared skull and cured with UV light. Dexamethasone (Patterson Veterinary, 4 mg/kg) was injected intraperitoneally to reduce swelling in the brain. A craniotomy was made over the cerebellum behind lambda for the ground electrode, which was secured to the skull using an optical dental composite. A second, larger craniotomy was made above the hippocampus (AP: −2.0 mm, ML: +1.5 mm from bregma) using a trephine stainless steel bur. Surgilube was applied to the surface of the brain, and the bundle of the drive was dipped in mineral oil. The drive was lowered until the bundle rested on the surface of the brain and secured to the skull using optical dental composite and UV light. The ground of the drive and the implanted ground wire were soldered together, and the excess ground wire was trimmed and encased in the dental composite to protect the ground wire and blunt any sharp edges. Mice were removed from the stereotaxic frame and carprofen was administered subcutaneously as an analgesic. Immediately after the surgery, each tetrode was turned down 450 µm into cortex.

### 2.4 Data acquisition during random foraging in an open field

Two weeks after drive implantation, single units and video were recorded simultaneously during all rest and behavioral sessions. Recordings were made daily for a duration of one week. Thus, data presented in this manuscript are obtained six to seven weeks after epilepsy induction. The electrophysiological data were synchronized with video using Bonsai (36, 37) and data were acquired using the Open Ephys Acquisition Board (37). Following a one-week postoperative recovery period, mice were food restricted to maintain 85% of their baseline weight. Mice were trained to randomly forage for food reward (Bio-Serv, Bacon Flavored Foraging Crumbles, #F5783) in an open, black, square arena (diameter = 41 cm, height = 36 cm), with a checkered cue card. Behavioral sessions included: a 20-minute rest session before the first foraging session, four 10-minute foraging sessions with 5 minutes of rest in between, and a 20-minute rest session after the last foraging session. During this training period, tetrodes were strategically advanced into the hippocampus, targeting subfields CA1, CA3, and DG.

Once mice covered the arena consistently during behavioral sessions (fully covering the arena twice in 10 minutes) and tetrodes were in their target location, mice were then exposed to a novel arena. The novel arena was an open, white circle arena (diameter = 49 cm, height = 33 cm), with a checkered cue card. The foraging session sequence began with one session in the familiar square arena, followed by four sessions in the novel, circle arena, and ended with the last session in the familiar arena (Figure 3A). Again, foraging sessions were flanked by 20-minute rest sessions.

### 2.5 Histology and DG dispersion scoring protocol

At the conclusion of the experiment, mice received an overdose of sodium pentobarbital. Mice were perfused intracardially with 0.37% sulfide solution, followed by 4% paraformaldehyde (PFA) in phosphate-buffered saline (PBS). Brains were extracted and stored in the PFA solution overnight and then transferred to 30% sucrose in PBS. Brains were sectioned using a freezing microtome. 50 µm coronal sections were collected through the hippocampal segment with electrode tracks. Each section was mounted onto microscope slides using a gelatin solution in PBS. Mounted sections were double stained with TIMM sulfide silver and cresyl violet to visualize tetrode tracks. The final position of each tetrode was reconstructed using serial sections. Data from tetrodes were only included in the data analysis if the tetrode’s final position was determined to be in CA1, CA3, or DG cell layers.

DG dispersion measurements were determined by an investigator that manually measured the thickness of the upper blade of the DG for three consecutive sections of dorsal hippocampus, ipsilateral to the injection, for saline control and KA mice (Figure S4). The thickness for the three sections was averaged to obtain a measure of DG dispersion for each mouse. Similarly, CA3 area was measured across the same three sections and averaged. Finally, a qualitative score was given for CA1; 2 for dense control-like CA1 layer, 1 for a patchy CA1 layer, and 0 for a completely sclerotic layer.

### 2.6 Interictal spike (IS) detection on local field potential (LFP)

All signal processing was done in MATLAB (MathWorks, R2022b). The IS detection procedure used was based on Yi et al. (2025) (30). An LFP channel was selected from a hippocampal tetrode located in the CA1 cell layer for each KA mouse, during a familiar behavioral foraging session. The LFP was down-sampled to 1 kHz and band-pass filtered. Spikes that had peaks with a minimum peak prominence and peak height above a tuned threshold were counted as IS. Thresholds were manually based on user input for each session and ranged from 500 – 1000 μV.

### 2.7 Data processing and place cell classification

Electrophysiological data from each rest and behavioral session were high-pass filtered, and background subtracted for clustering into putative single units using Kilosort (38). Putative single units were then manually curated in Phy (https://github.com/cortex-lab/phy), where any cluster that did not show stable waveform amplitudes or clear refractory periods were discarded. The position of mice was estimated using DeepLabCut (39, 40) and correlated with synchronized spike times. Using position, instantaneous running speeds for each session were calculated. Position data and their respective spike times from periods of immobility, defined as a running speed less than 2 cm/s, were excluded.

Putative principal neurons were distinguished from putative interneurons using a 10 Hz average firing rate criterion (41, 42). Principal neurons were then categorized as either active or inactive using a peak firing rate criterion of 2 Hz. Inactive neurons (peak firing rate < 2 Hz) were excluded from the analysis. Active, putative principal neurons then underwent a shuffling procedure to classify them as either place or non-place cells. First, spatial information for each behavioral session was calculated using the following equation (43, 44):

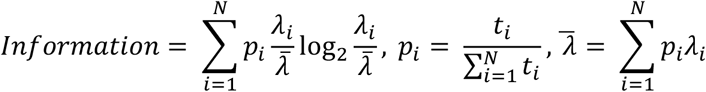

where i represents the 2 cm bins of the environment, p_i_ is the occupancy probability and λ_i_ is the mean firing rate of a given bin i, and λ is the overall mean firing rate of a cell. The session with the highest spatial information for a given cell was selected and then underwent a circular shuffling procedure (1000 shuffles in which spatial positions were time-shifted by a random amount and spatial information was recalculated). Putative principal neurons were classified as place cells if the if the original distribution was greater than the 95th percentile of the shuffled distribution. In addition, burst indexes were calculated for individual cells by dividing the number of spikes that occurred within 5 ms of one another by the number of spikes that occurred within 250 ms of one another.

### 2.8 Spatial analysis

#### Rate maps

Rate maps of neuronal firing for each behavioral session were constructed as described in Skaggs et al. (1996) (44). Environments were divided into 2 cm bins, and the firing rate at each location was calculated by expanding a circle centered on a given bin until it met the following criterion:

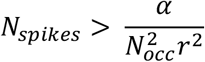

where N_spikes_ is the number of spikes, N_occ_ is the number of occupancy samples with a sampling frequency of 30 Hz, r is the radius of the circle in bins, and α is a scaling factor set as 10,000. The firing rate at each location was calculated as the following:

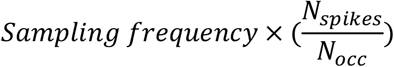

#### Spatial correlations

To assess spatial stability within a session, rate maps were split into the first and second half of the session. The correlation coefficients (r) between half maps were calculated using Pearson’s correlation. To assess spatial stability across sessions, correlation coefficients were calculated between two behavioral sessions using Pearson’s correlation.

#### Spatial coherence

A vector of the rate map was constructed that preserves neighboring spatial bin relationships and the autocorrelation of that vector was calculated using the *corrcoef* Matlab function.

#### Sparsity

Calculated using the following equation (43, 44):

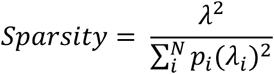

#### Spatial selectivity

Peak firing rate (in the spatial bin with the highest rate) divided by the average firing rate calculated across all spatial bins. p_i_ is the occupancy probability and λ_i_ is the mean firing rate of a given bin i, and λ is the overall mean firing rate of a cell.

#### Place field boundaries

Individual spatial firing fields of neurons were isolated as previously described (45). Fields were calculated on the session with the highest spatial information of each place cell. For each firing peak in the rate map (>0.5 Hz), contours were calculated at 20 levels between zero and the peak rate. For overlapping contours between fields, each shared contour was divided into segments at inflection points of the contour, and each segment was assigned to the nearest field. After fields were determined, only fields with peak rates >2 Hz and that had areas of at least 20 cm^2^ were included for further analysis.

### 2.9 Statistical analysis

Statistics were performed using MATLAB (MathWorks, R2022b) software. To compare cell properties and theta coherence between experimental groups, we used a generalized linear mixed model design (*fitglme* in MATLAB). Mouse and cells nested within mouse (when applicable) were included as random factors in the model. Experimental group (i.e., control or KA) was set as a fixed factor. In additional to p-values, the t-statistic associated with each mixed effect model for the experimental group coefficient is reported. When applicable, the Bonferroni correction was used for multiple comparisons. For by animal measures, Pearson’s correlations were performed on mean values from individual animals.

## 3 RESULTS

### 3.1 Proportion of place cells are reduced in hippocampal subfield CA1 in KA mice

Our study sought to explore the impact of chronic, focal TLE on hippocampal function. Here, we investigated spatial coding properties of the three hippocampal subfields (CA1, CA3, and DG) in epileptic mice, ipsilateral to the epileptic focus. We employed the supra-hippocampal kainate model of TLE (31, 46, 47). Mice were injected into the cortex above hippocampus with kainic acid (KA; N = 10, 6 male/4 female) or saline as a control (N = 6, 3 male/3 female). Four weeks post-injection, mice were implanted with an Open Ephys ShuttleDrive (35) and electrophysiological data from the three hippocampal subfields were obtained while mice foraged in a familiar environment (Figure 1A).

**Figure 1.**
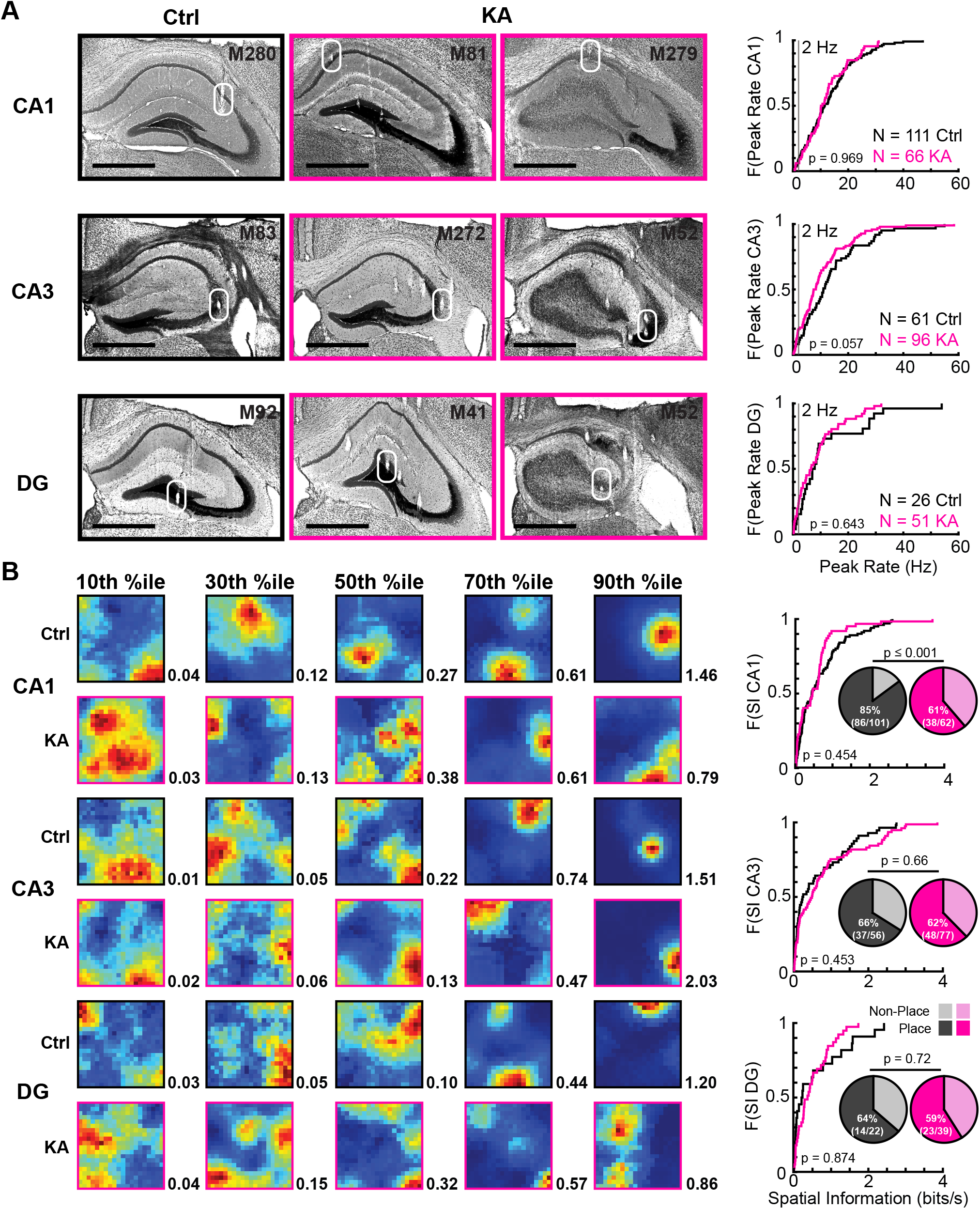
Reduced number of place cells in area CA1 of KA mice. ***A left***, Coronal sections through the hippocampus with tetrode tracks from control (black) and KA (pink) mice. The tetrode positions for recordings are indicated with a white circle for hippocampal CA1, CA3, and DG subregions. Scale bar, 870 µm. ***A right***, Cumulative distribution functions for spatial peak firing rates of putative principal neurons in CA1, CA3, and DG of control and KA mice. Grey line at 2 Hz shows cutoff for defining principal neurons as ‘active’. No significant differences in peak spatial firing rates were found between groups for any sub-region (GLME, see supplemental material for parameters and statistics). ***B left***, Hippocampal putative principal cells recorded from CA1, CA3, and DG organized by spatial information for control and KA mice. Spatial information was calculated for putative principal neurons that were active during foraging. Example rate maps are shown, organized from low to high spatial information. Spatial information (black) is indicated to the right of each example map. ***B right***, Cumulative distribution functions for spatial firing rates for all active neurons recorded in CA1, CA3, and DG of control and KA mice. No significant differences were found between groups for any sub-region (GLME, see supplemental material for parameters and statistics).The proportion of place and non-place cells in CA1, CA3, and DG for control (dark/light grey) and KA (dark/light pink) mice are represented by pie charts and are inset in each cumulative distribution plot. Single units were categorized as place cells if their spatial information was greater than the 95th percentile of a shuffled distribution (see methods). The proportion of place cells in CA1 of KA mice was significantly smaller than in controls (χ^2^ = 12.01, *p* ≤ 0.001). Significance denoted with asterisks: ***p ≤ 0.001.

We obtained recordings from neurons from each hippocampal subfield while mice foraged in familiar spatial environments. Mean rates and burst indexes (proportion of spikes occurring within 5 ms) were calculated for all neuron recorded (Figure S1). Putative principal neurons were separated from putative interneurons based on mean firing rates. For each putative principal neuron, spiking during foraging was spatially binned and spatial peak rates were calculated to determine neurons that were active during foraging behavior. There was no significant difference in the distributions of spatial peak firing rates between control and KA mice in any subregion (Figure 1A). Next, we calculated spatial information for each neuron - a measure of how well a given spike from a neuron would predict the animals location in the environment (Figure 1B), and found no significant difference between subgroups in any region. Using the spatial information measure, we next classified cells as place cells if they had spatial information that was significantly higher than chance levels determined by a shuffling procedure for each cell (Figure 1B, inset pie charts, *see methods*). We found that the proportion of place cells was significantly lower in CA1 of KA mice compared to control mice (Figure 1E, χ^2^(1,163) = 12.01, *p* ≤ 0.001, Chi-squared test). In contrast, there was no significant difference in the proportion of place cells between control and KA mice for CA3 (Figure 1E, χ^2^(1,133) = 0.20, *p* = 0.66, Chi-squared test) or DG (Figure 1E, χ^2^(1,61) = 0.13, *p* = 0.72, Chi-squared test). Taken together, despite similar activity levels in KA and control mice, we observe general place coding deficits in CA1.

### 3.2 Region specific impairments of spatial coding in familiar environments are impaired in KA mice

Next, we assessed the quality of place coding in familiar environments (Figure 2, Table 1 for cell breakdown by region and animal). For each parameter tested, we observed impairments in at most one sub-region. In CA3, place cells exhibited less within session stability in KA mice compared to control mice (Figure 2B). In CA1, place cells exhibited less spatial coherence in KA mice compared to control mice (Figure 2C). Finally, in DG, place cells had lower spatial selectivity in KA mice compared to in control mice (Table 2). Both spatial coherence and selectivity would be impacted if place cells comprised more place fields, however we found no difference in number of fields or field size (Figure S2). Together, these data point towards subregion-specific impairments in place coding across the hippocampal circuit in KA mice.

**Table 1.**
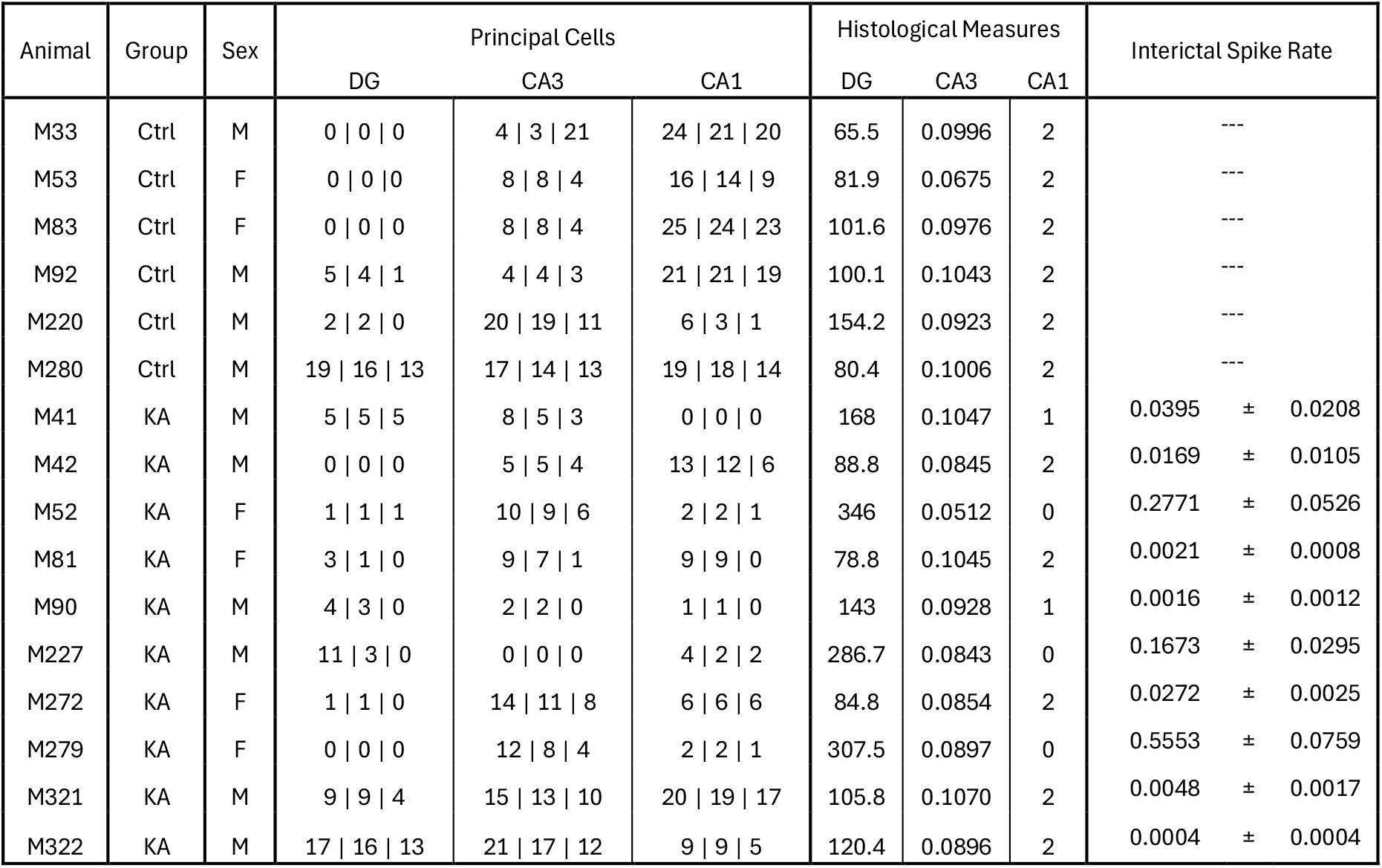
**Animal:** mouse ID. **Group:** Ctrl, control saline injected; KA, KA injected. **Sex:** M, male; F, female. **Principal Cells:** total number recorded | total number that were active in at least one behavioral session (> 2 Hz peak rate) | total number of place cells (significantly spatially informative after a shuffling procedure). **Histological Measures:** DG, dentate gyrus cell layer width measurement in μm; CA3 cell layer area in mm^2^; CA1, categorical sclerosis score, 2 no sclerosis, 1 intermediate sclerosis, 0 complete sclerosis. **Interictal Spike Rate:** Average interictal spike rate calculated across all foraging sessions (Hz).

| Animal | Group | Sex | Principal Cells |  |  | Histological Measures |  |  | Interictal Spike Rate |
| --- | --- | --- | --- | --- | --- | --- | --- | --- | --- |
|  |  |  | DG | CA3 | CA1 | DG | CA3 | CA1 |  |
| M33 | Ctrl | M | 0 \| 0 \| 0 | 4 \| 3 \| 21 | 24 \| 21 \| 20 | 65.5 | 0.0996 | 2 | --- |
| M53 | Ctrl | F | 0 \| 0 \| 0 | 8 \| 8 \| 4 | 16 \| 14 \| 9 | 81.9 | 0.0675 | 2 | --- |
| M83 | Ctrl | F | 0 \| 0 \| 0 | 8 \| 8 \| 4 | 25 \| 24 \| 23 | 101.6 | 0.0976 | 2 | --- |
| M92 | Ctrl | M | 5 \| 4 \| 1 | 4 \| 4 \| 3 | 21 \| 21 \| 19 | 100.1 | 0.1043 | 2 | --- |
| M220 | Ctrl | M | 2 \| 2 \| 0 | 20 \| 19 \| 11 | 6 \| 3 \| 1 | 154.2 | 0.0923 | 2 | --- |
| M280 | Ctrl | M | 19 \| 16 \| 13 | 17 \| 14 \| 13 | 19 \| 18 \| 14 | 80.4 | 0.1006 | 2 | --- |
| M41 | KA | M | 5 \| 5 \| 5 | 8 \| 5 \| 3 | 0 \| 0 \| 0 | 168 | 0.1047 | 1 | 0.0395 ± 0.0208 |
| M42 | KA | M | 0 \| 0 \| 0 | 5 \| 5 \| 4 | 13 \| 12 \| 6 | 88.8 | 0.0845 | 2 | 0.0169 ± 0.0105 |
| M52 | KA | F | 1 \| 1 \| 1 | 10 \| 9 \| 6 | 2 \| 2 \| 1 | 346 | 0.0512 | 0 | 0.2771 ± 0.0526 |
| M81 | KA | F | 3 \| 1 \| 0 | 9 \| 7 \| 1 | 9 \| 9 \| 0 | 78.8 | 0.1045 | 2 | 0.0021 ± 0.0008 |
| M90 | KA | M | 4 \| 3 \| 0 | 2 \| 2 \| 0 | 1 \| 1 \| 0 | 143 | 0.0928 | 1 | 0.0016 ± 0.0012 |
| M227 | KA | M | 11 \| 3 \| 0 | 0 \| 0 \| 0 | 4 \| 2 \| 2 | 286.7 | 0.0843 | 0 | 0.1673 ± 0.0295 |
| M272 | KA | F | 1 \| 1 \| 0 | 14 \| 11 \| 8 | 6 \| 6 \| 6 | 84.8 | 0.0854 | 2 | 0.0272 ± 0.0025 |
| M279 | KA | F | 0 \| 0 \| 0 | 12 \| 8 \| 4 | 2 \| 2 \| 1 | 307.5 | 0.0897 | 0 | 0.5553 ± 0.0759 |
| M321 | KA | M | 9 \| 9 \| 4 | 15 \| 13 \| 10 | 20 \| 19 \| 17 | 105.8 | 0.1070 | 2 | 0.0048 ± 0.0017 |
| M322 | KA | M | 17 \| 16 \| 13 | 21 \| 17 \| 12 | 9 \| 9 \| 5 | 120.4 | 0.0896 | 2 | 0.0004 ± 0.0004 |

**Table 2.**
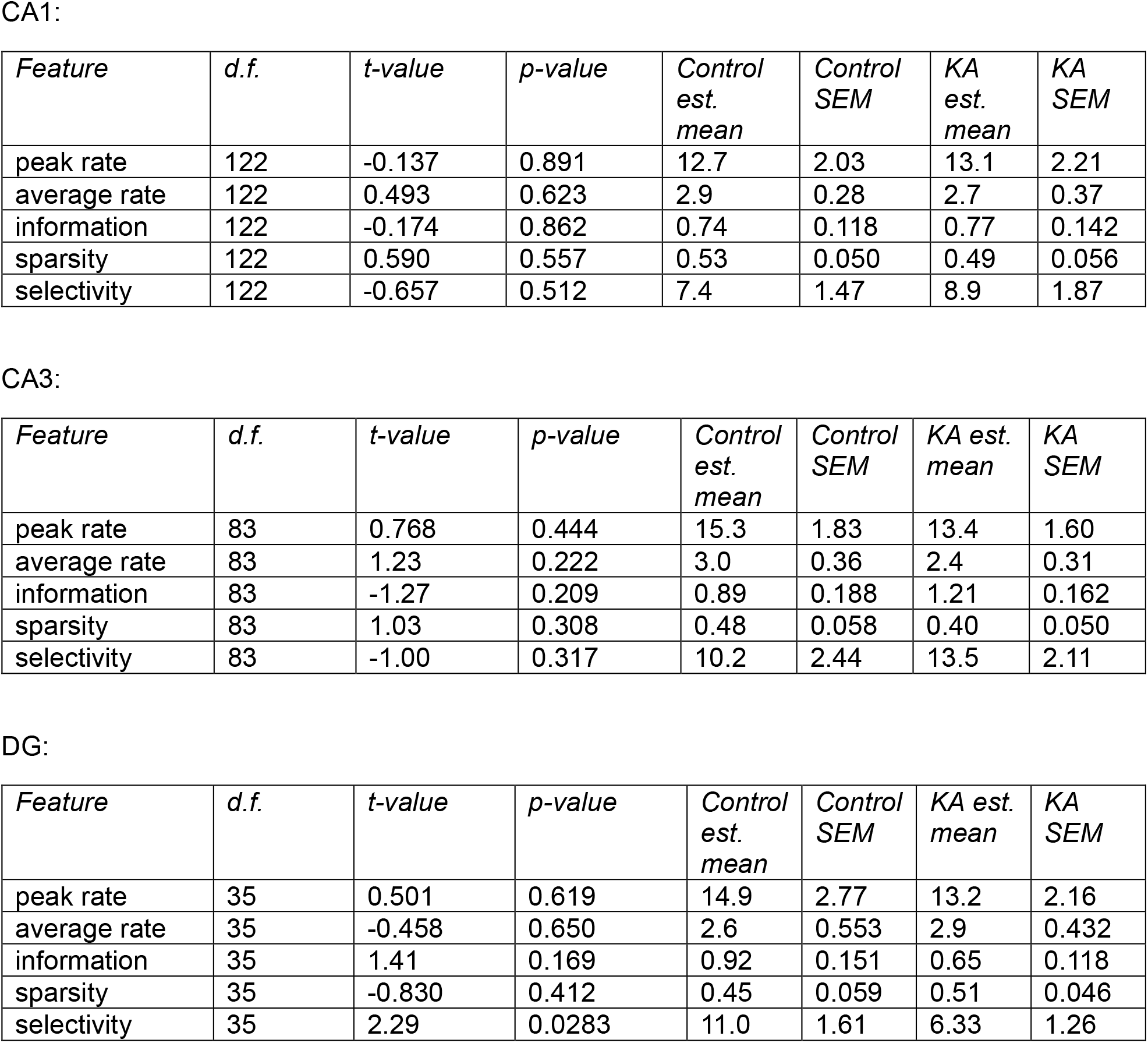
Additional properties of place cells for hippocampal CA1, CA3, and DG. For each hippocampal region and feature, a GLME was performed on the estimated mean.

**Figure 2.**
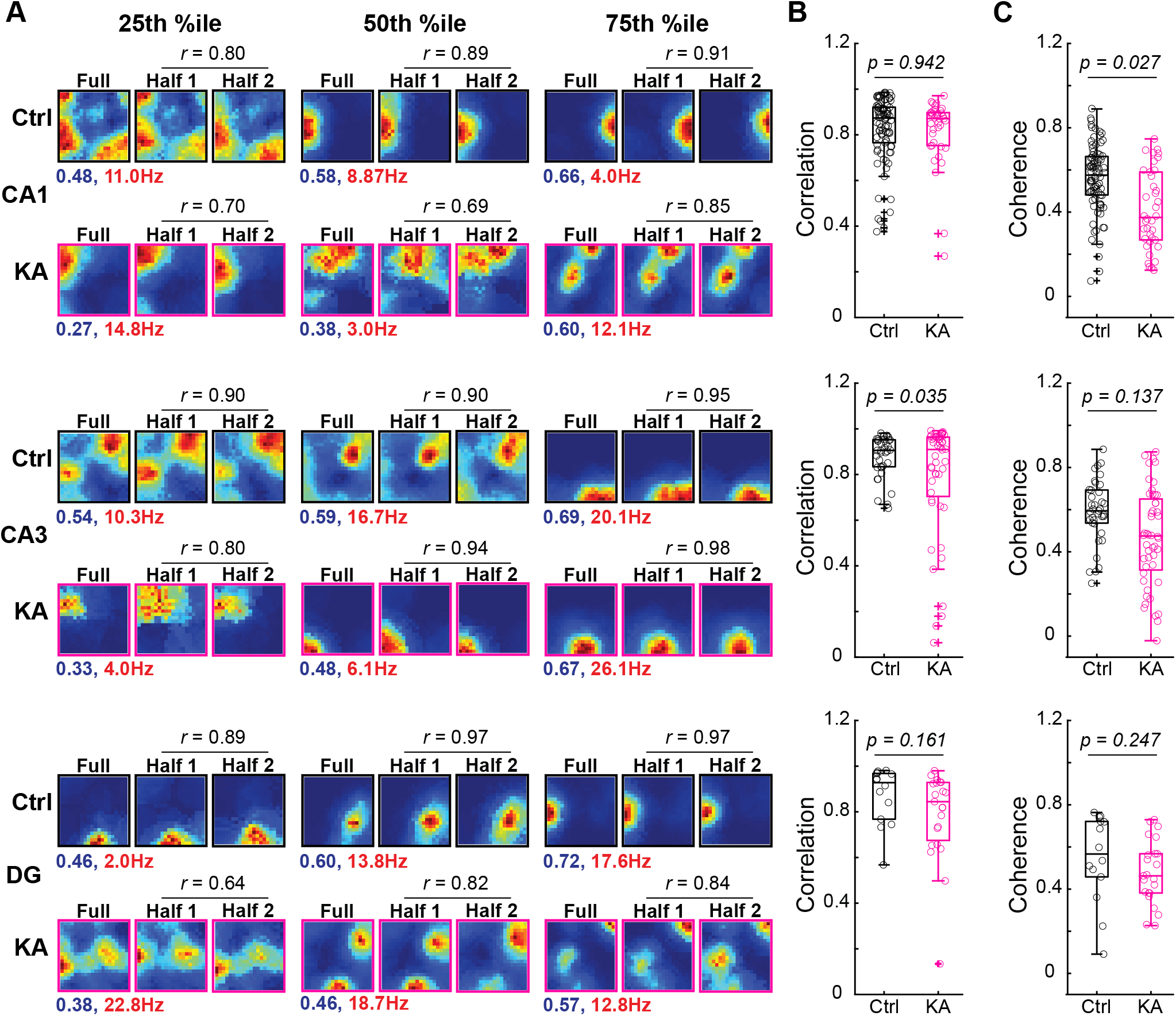
Region specific disruption of spatial parameters of place cells in KA mice. ***A***, Example rate maps of place cells from CA1, CA3, and DG for control (black) and KA (pink) mice. Example maps shown are the 25^th^, 50^th^, and 75^th^ percentile for spatial coherence. For each example, the full map and two half maps (first and second half of the recording) are shown. The spatial coherence (blue), spatial peak firing rate (Hz, red), and spatial correlation (black) are indicated for each example map. ***B***, Within session spatial correlations (calculated from half maps) of place cells from CA1, CA3, and DG for control (black) and KA (pink) mice. Box-and-whisker plots of correlations with the correlations for individual cells plotted as open circles. Median correlations are indicated with a solid line. Within session correlation was significantly lower for CA3 of KA mice compared to control (GLME, t(83) = 2.1, p = 0.0350; estimated means: control = 0.88 +/−0.032; KA = 0.79 +/−0.0281). ***C***, Spatial coherence of place cells from CA1, CA3, and DG from control and KA mice. Box-and-whisker plots of spatial coherence with coherences for individual cells plotted as open circles. Median coherences are indicated with a solid line. Spatial coherence was significantly lower in CA1 of KA mice compared to control (GLME, t(122) = 2.24, p = 0.027; estimated means: control = 0.55 +/−0.040, KA = 0.42 +/−0.044). See supplemental material for full statistical parameters.

An important consideration is that several aspects of epilepsy emerge at the mouse level, rather than at the cellular level. It is important to assess such effects to understand the network landscape that place coding changes occur in (note that the GLMEs account for animal identity as a random factor). First, we tested whether there were differences in oscillatory coherence across subregions and found no difference between control and KA mice in all of the oscillatory bands tested (Figure S3). We noted that across KA mice, we observed substantial variability in rates of interictal spikes and measures of pathohistology. There was a correlation between interictal spike rate and average spatial information in CA1 (Figure S4), however measures of place maps such as correlation and coherence did not correlate with interictal spike rate in any region tested (DG was not included due to low numbers of animals with DG place cells and non-zero interictal rates, Table 1). Although we observed variable histopathology, the extent of pathology between subfields was correlated across animals (Figure S5A, B). In other words, animals with large granule cell dispersion also exhibited a sclerotic CA3, thus we focused our correlation analysis on granule cell dispersion which was the parameter we felt we captured most reliably. There was no correlation between spatial coding properties and the extent of dispersion – suggesting that place field problems arise in networks that appear more anatomically intact.

### 3.3 Spatial stability of newly formed maps in a novel environment are impaired in KA mice

We next examined another feature of hippocampal place cells, the ability to quickly establish new representations (i.e., place maps) upon entering a completely novel environment (Figure 3). Electrophysiological recordings were obtained from the three hippocampal subfields (CA1, CA3, and DG) while mice explored both a familiar and a novel environment (Figure 3A-B). We observed that all subfields of the hippocampus remapped between familiar and novel environments (i.e., formed unique spatial representations) for control mice and KA mice. Additionally, the extent of remapping was of similar magnitude between control and KA mice for all subregions (Figure 3D, CA1: z = −0.90, *p* = 0.37; CA3: z = −0.80, *p* = 0.42; DG: z = 0.45, *p* = 0.75, Wilcoxon signed rank test). Next, we assessed the stability of the newly formed maps across multiple exposures to the same novel environment (Figure 3E). We observed a trend toward decreased stability of newly formed maps in CA3 of KA mice (Figure 3E, p = 0.062). Finally, there was no significant difference in the stability of the newly formed place maps between control and KA mice for CA1or DG (Figure 3E). These findings demonstrate that remapping is intact in KA mice across the major hippocampal subregions.

**Figure 3.**
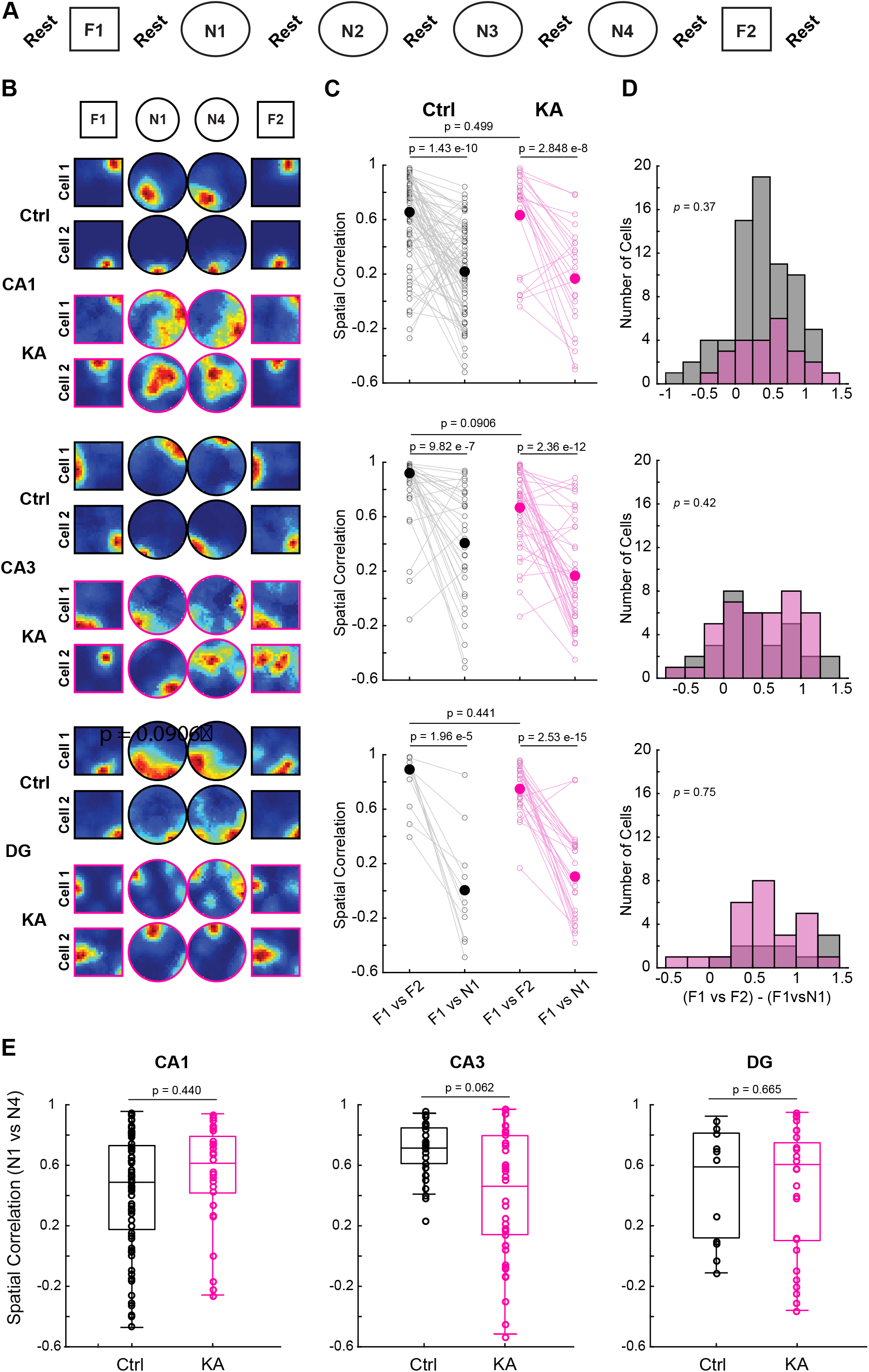
Hippocampus is able to generate new maps in mice with epilepsy. ***A***, Schematic of the novel room experimental paradigm. The experiment began with a 20-minute rest session before the first foraging session, followed by six 10-minute foraging sessions with 5 minutes of rest in between. The experiment concluded with a 20-minute rest session after the last foraging session. The foraging session sequence began with one session in the familiar arena (F1), followed by four sessions in the novel arena (N1-N4), and ended with one session back in the same familiar arena (F2). ***B***, Example heat maps for individual place cells from hippocampal subregions CA1, CA3, and DG for control (black) and KA (pink) mice. ***C***, Spatial correlations for comparisons between F1-F2 and F1-N1 for hippocampal CA1, CA3, and DG from control and KA mice. The median spatial correlation for each comparison is plotted as solid circles with spatial correlations for each individual cell plotted as paired, open circles. All statistics presented are on estimated means from GLMEs (see supplement for full statistical parameters). ***D***, Distribution of place cell spatial correlations for the difference between F1-F2 and F1-N1. There was no significant difference between distributions of control and KA mice in CA1 (z = −0.90, *p* = 0.37), CA3 (z = −0.80, *p* = 0.42), or DG (z = 0.45, *p* = 0.75), Wilcoxon rank sum test. ***E***, Spatial correlations of place cells from comparisons of N1-N4 for hippocampal CA1, CA3, and DG from control and KA mice. Box-and-whisker plots are shown for each hippocampal subregion for control and KA mice. The median spatial correlations are indicated as a solid line. All statistics presented are on estimated means from GLMEs (see supplement for full statistical parameters).

## 4 DISCUSSION

We show unique alterations in place representations of hippocampal subfields in a rodent model of focal temporal lobe epilepsy (TLE). In familiar environments, there were fewer place cells in CA1 of epileptic mice and those remaining exhibited a reduction in spatial coherence. Our findings are consistent with results from previous studies of CA1 place coding in TLE with respect to reduced map quality (4, 5, 6, 7, 8, 9), but differ in that we did not observe reduced stability of place fields in CA1. The majority of studies of place coding in CA1 have been done using systemic administration of excitotoxic drugs to induce epilepsy. In those cases, it is likely that the changes to the temporal lobe are more extensive (involving bilateral and ventral temporal lobe structures at the very least). Thus it is possible that in our focal model, CA1 is still receiving relatively normal inputs from contralateral CA3 etc. which allows for stabilization of fields. It is interesting to speculate that a driver of fewer cells exhibiting significant spatial information in CA1 is interictal spike rates (Figure S4). An interesting follow up study would be to see whether inhibition of interictal spiking would rescue the number of place cells in the CA1 subfield.

In CA3 place cells, we observed reduced stability within familiar environments and a trend that new maps formed in novel environments were less stable across sessions. A recent study suggests that map stability in CA3 neurons is correlated with the neurons position within the proximal distal axis of CA3 – which also correlates with the amount of recurrence (48). Less stability was observed in the cells with stronger recurrence (reflecting updating of autoassociative network across experience). Thus, it would be interesting to test whether in TLE, there is stronger recurrence, either via improperly strengthened synapses (as has been demonstrated for the input to CA3 from DG (49)) or sprouting of recurrent axons (50).

In DG place cells, we observed a reduction in selectivity. Spatial selectivity is essentially a measure of signal to noise of a place map. Given that peak rates and field numbers were similar between groups – it suggests that out-of-field firing is slightly elevated in DG cells. Such an effect is in line with impaired feedback inhibition. Loss of inhibition within the hippocampal circuit is widely reported in TLE, and others have shown that disrupted inhibition in DG/CA3 impacts place coding in CA1 and DG (51, 52, 53). Importantly, we do not distinguish between granule cells and mossy cells in this study.

Another potential contributor to the disparate deficits we observe is the upstream input coming from the entorhinal cortex, which differentially innervates the DG/CA3 and CA1 subfields via the perforant pathway and temporo-ammonic pathway, respectively (54, 55). Along these lines, lesions of medial entorhinal cortex (MEC) layer 3 cause a reduction in spatial coherence of CA1 place cells (56), and impairments in excitatory neurons from MEC layer 3 have been reported in TLE (57). Multiple studies have shown a reduction in inhibitory synaptic input of MEC layer 2 neurons that innervate DG and CA3, resulting in increased synaptic input to these regions (25, 58, 59).

Past TLE research focused almost exclusively on place coding in a familiar environment or similar familiar environments (53), with little known about how the development and construction of new maps in novel environments would be impacted. We found that while control and KA mice formed distinct spatial representations between familiar and novel environments across hippocampal subregions, there was a trend toward impaired spatial stability of the newly formed place maps in CA3 of KA mice. Seminal research on remapping in the hippocampus has shown that distinct spatial maps are formed in novel environments but stabilized at different rates in the main hippocampal subregions of healthy, control animals (10, 60). Specifically, newly formed place maps from CA3 take longer to stabilize than CA1 when introduced to a novel environment – and our findings suggest this stabilization is exacerbated in epilepsy, which again may point to augmentation of recurrence in CA3 (61). Novelty has been shown to drive DG granule cells and switch downstream CA3 networks into a state of forming new memories (i.e., spatial representations) rather than recalling previously formed maps (62). In TLE, this switch could be altered by the loss of inhibition from DG, resulting in a stronger drive to CA3 to form new maps and thus a destabilization across multiple exposures to the same novel environment.

## 5 CONCLUSIONS

In the epileptic brain, we found problems with map quality in CA1; there was a decrease in the proportion of place cells place fields of CA1 were less coherent. We found problems with stability in CA3 cells and problems with selectivity in DG. To further assess mechanisms of remapping and stability of newly formed place maps, we performed recordings while mice explored a novel environment and found that all subregions constructed new, distinct maps within the first session of exposure to a novel environment, however new maps in CA3 trended toward being less stable in epileptic mice. Together, our findings highlight that focal TLE causes impairments in the spatial codes of place cells across hippocampal subfields CA1, CA3, and DG. Problems in CA1 could be partially explained by interictal spike rates during foraging, but no other feature of place coding problems correlated with interictal spike rates, nor could any place coding problems be explained by anatomical changes. Subregion-specific impairments demonstrate how TLE impacts critical memory circuits and provide a lens into how these impairments could disrupt memory flexibility and cognition.

## ACKNOWLEDGEMENTS

NIH R01 1R01NS128222 (to L.A.E.), American Epilepsy Society Grant 835029 (to L.A.E.), American Epilepsy Society Grant 1462229 (to B.L.B.), and National Institute of Neurological Disorders and Stroke Grant T32 5T32NS045540 (to B.L.B.).

## AUTHOR CONTRIBUTIONS

BLB: Conceptualization (equal); Methodology (equal); Investigation (lead); Data Curation (lead); Formal Analysis (lead); Software (equal); Visualization (lead); Funding Acquisition (supporting); Supervision (equal); Writing – Original Draft Preparation (equal); Writing – Review and Editing (equal). MMD: Formal Analysis (supporting); Visualization (supporting) CD: Investigation (supporting). GT: Formal Analysis (supporting); Software (equal); Visualization (supporting). LAE: Conceptualization (equal); Methodology (equal); Formal Analysis (supporting); Software (equal); Visualization (supporting); Funding Acquisition (lead); Resources (lead); Supervision (equal); Writing – Original Draft Preparation (equal); Writing – Review and Editing (equal). All authors contributed to and approved the final manuscript.

