## Supplemental materials for "Unique deficits in place coding across subfields of the hippocampus in a mouse model of temporal lobe epilepsy"

These supplemental materials comprise two parts.

Part 1: Figures S1 – S5 and their corresponding figure legends.

Part 2: A statistical companion to the main text, in which we report the details of all of the statistics used. The statistical test, parameters, statistics, p-values, n, etc. are listed.

Figure S1. Boublil et al.

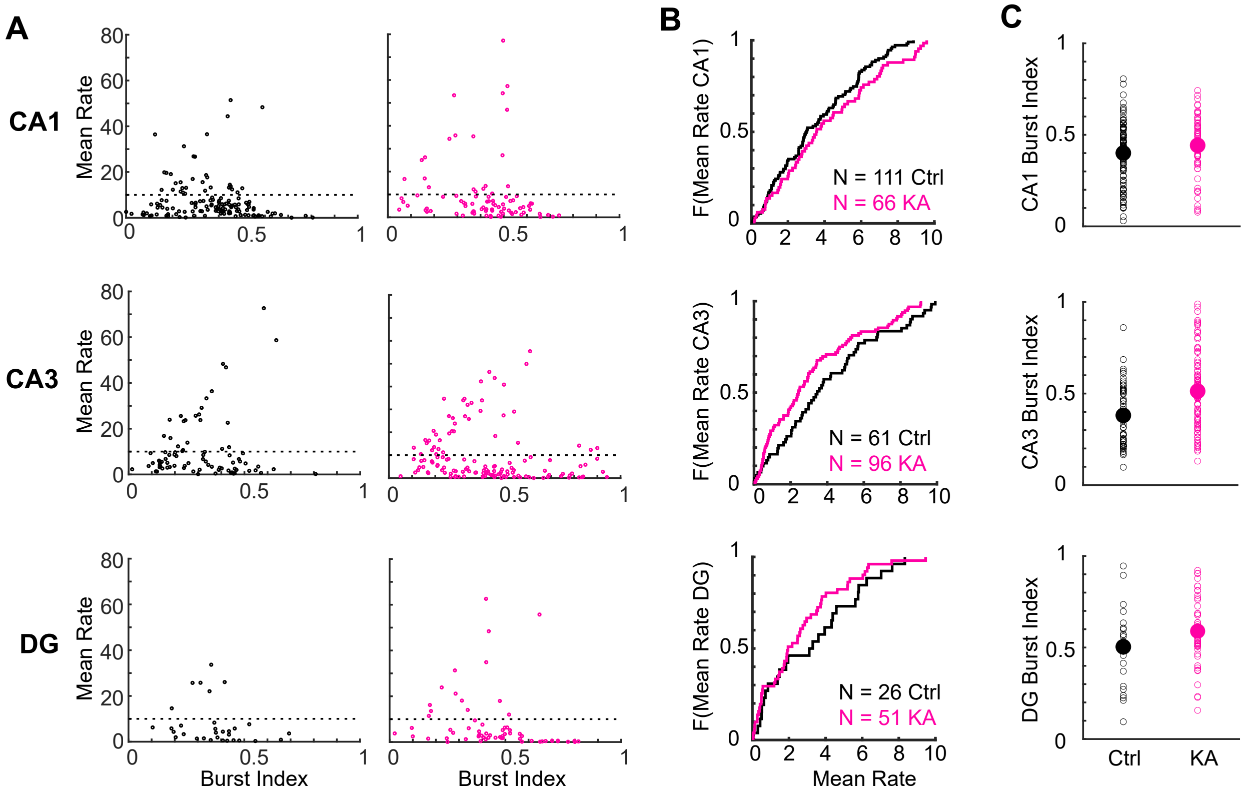

Figure S2. Boublil et al.

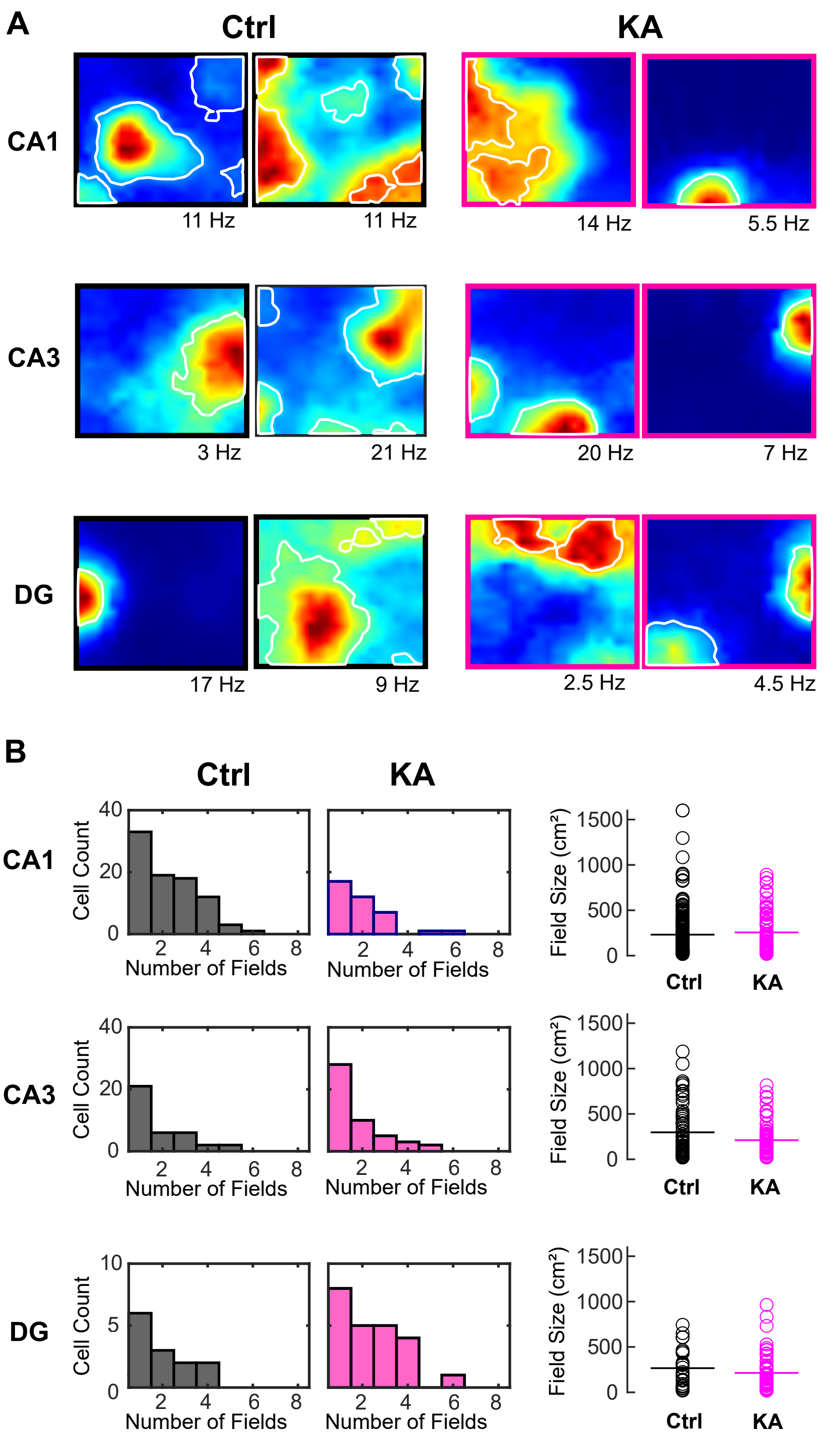

Figure S3. Boublil et al.

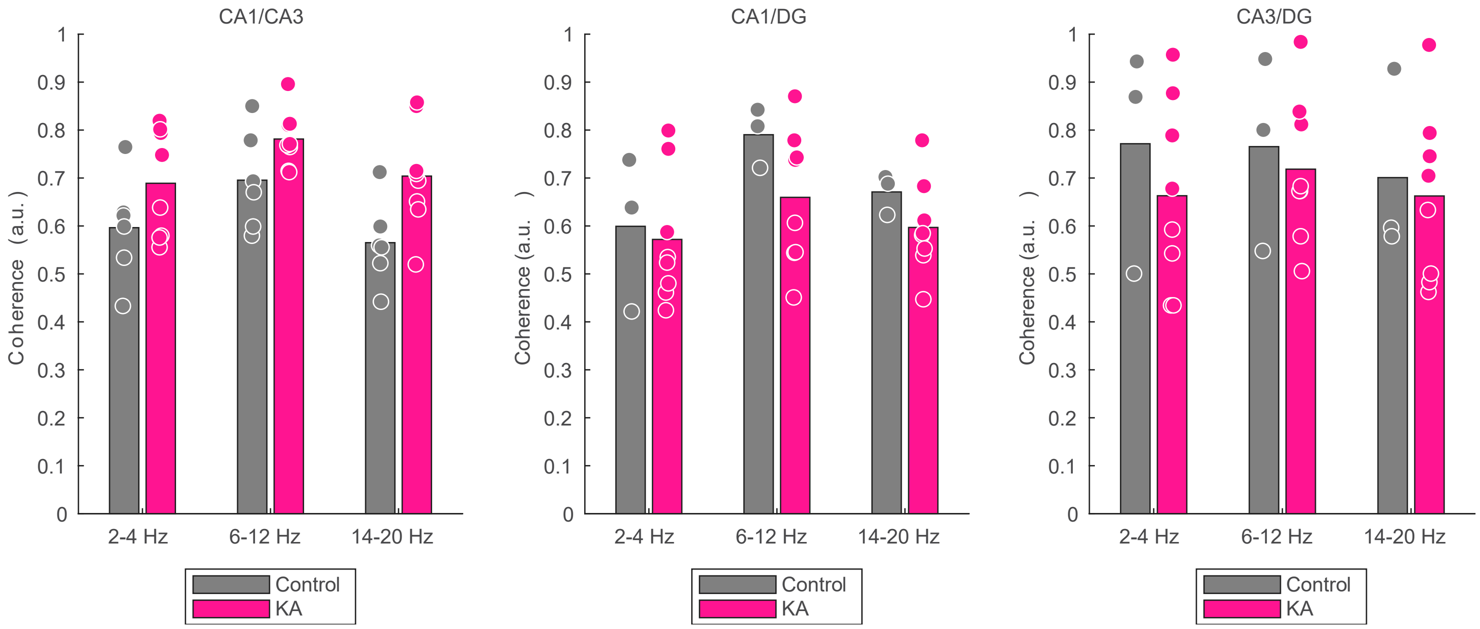

Figure S4. Boublil et al.

A

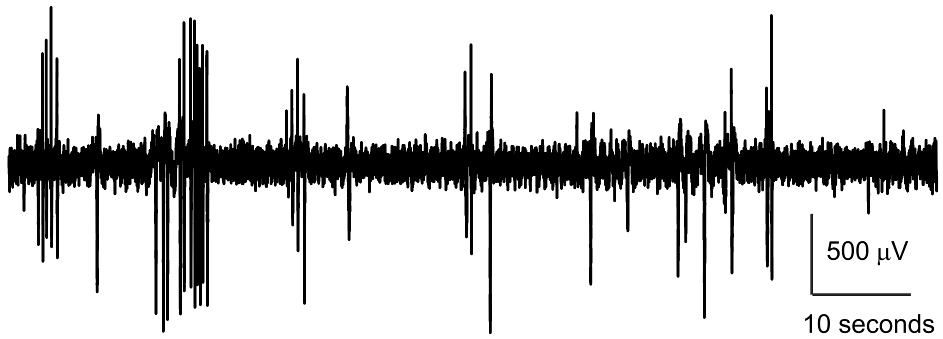


B

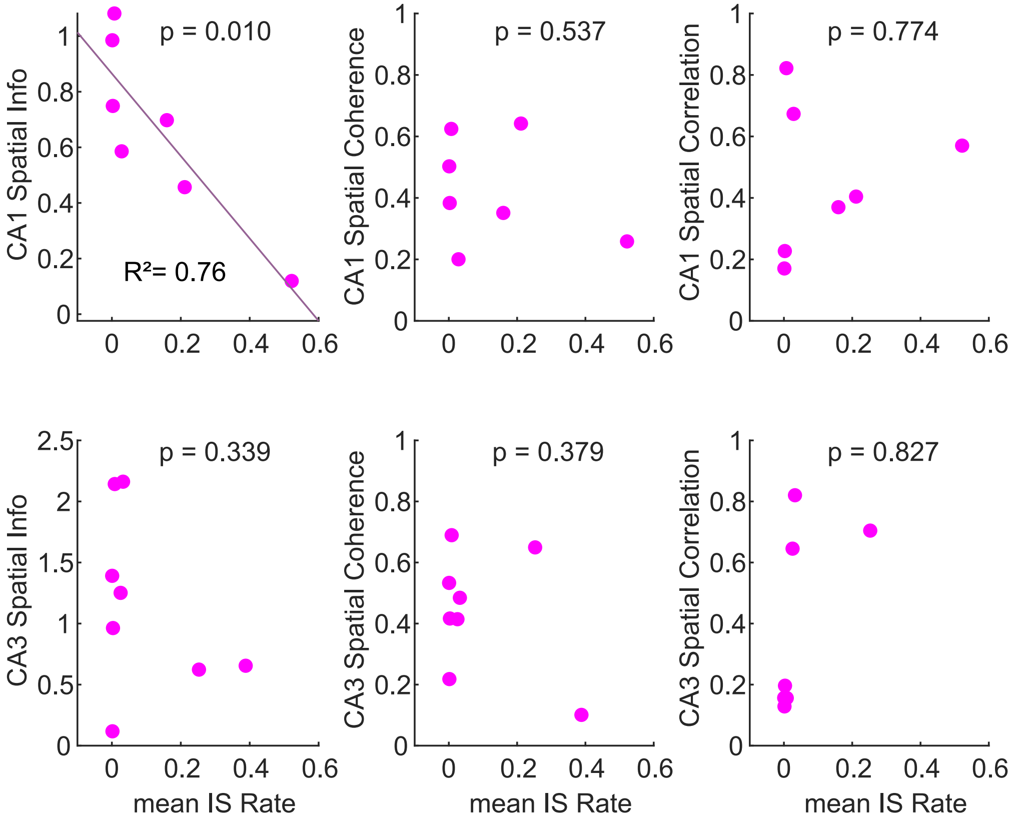

Figure S5. Boublil et al.

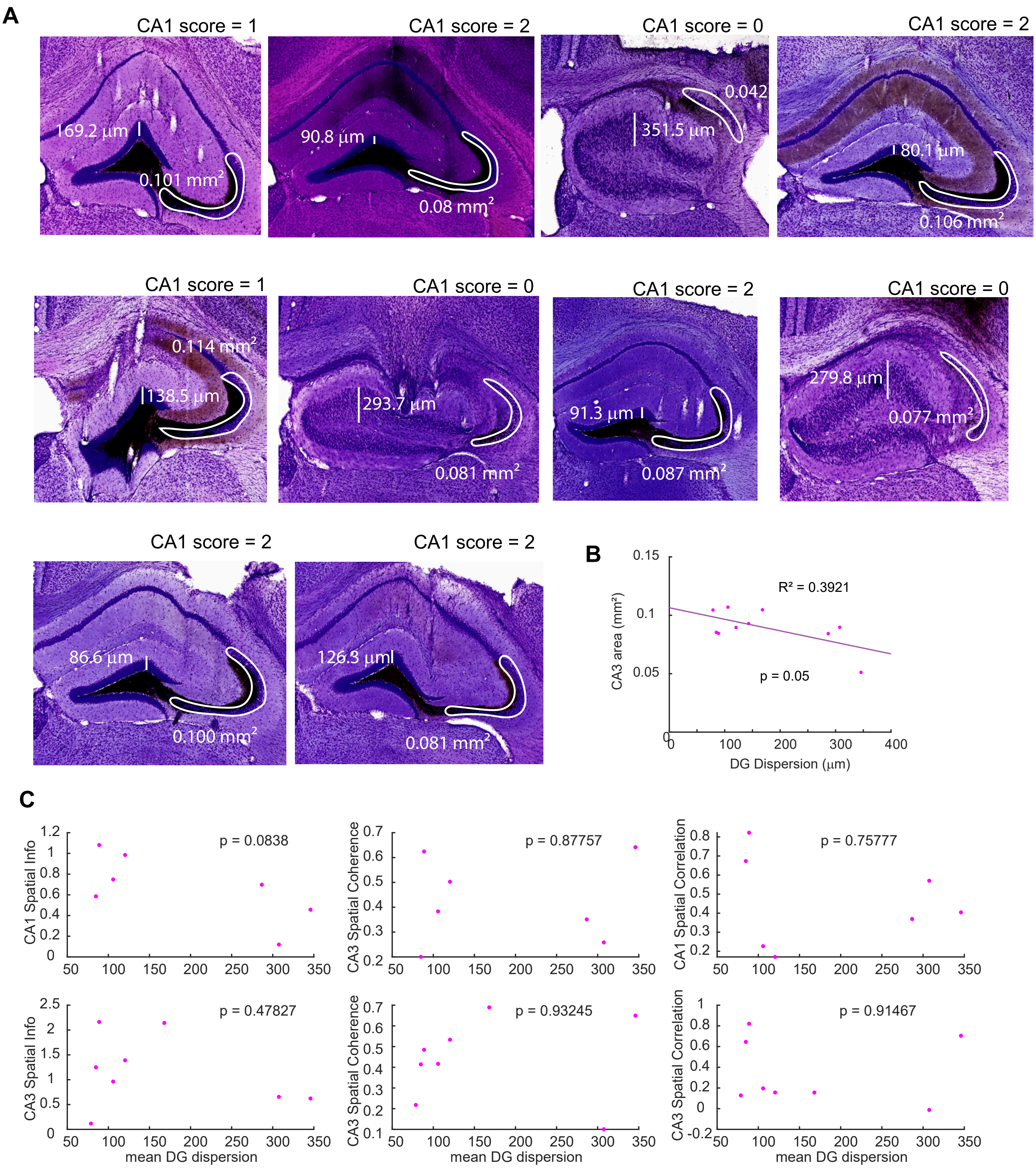

**Figure S1. Classification of putative principal neurons and interneurons. A,** Mean rate plotted as a function of burst index for all neurons recorded in CA1, CA3, and DG of KA and control mice. The dotted line shows the threshold at a mean rate of 10 Hz that was used to delineate between putative principal neurons and putative interneurons. **B,** Cumulative distributions of mean rates for all putative principal neurons. Generalized linear mixed effects model testing the effect of experimental group on max average firing rate for principal cells, tested for each region separately. No significant effect of experimental group for CA1 principal cell firing rate (control vs. KA, t(175) = -1.0, p = 0.319; estimated means: control = 3.38 +/- 0.585, KA = 4.20 +/- 0.561). Significant effect of experimental group for CA3 (control vs. KA, t(155) = 2.01, p = 0.0457; estimated means: control = 4.02 +/- 0.348, KA = 3.12 +/- 0.277. No significant effect of experimental group for DG (control vs. KA t(75) = -0.446, p = 0.657; estimated means: control = 3.09 +/- 0.481, KA = 2.70 +/- 0.740. **C,** Distributions of burst index for all putative principal neurons. Individual neurons are shown as open circles. Means are shown as large filled circles. Generalized linear mixed effects model testing the effect of experimental group on burst index for principal cells, tested for each region separately. No significant effect of experimental group for CA1 principal cell firing rate (control vs. KA t(173) = -0.522, p = 0.603; estimated means: control = 0.35 +/- 0.053, KA = 0.39 +/- 0.048). No significant effect of experimental group for CA3 principal cell firing rate (control vs. KA t(151) = -1.84, p = 0.0683). No significant effect of experimental group for DG principal cell firing rate (control vs. KA t(74) = 0.55, p = 0.585; estimated means: control = 0.32 +/- 0.050, KA = 0.44 +/- 0.040).

**Figure S2. Place field properties of place cells in KA and control mice. A,** Example heat maps of place cells with field bounds shown in white. Spatial peak rates (Hz) are shown below each map in black. All fields had spatial peak rates of at least 2 Hz. **B,** Left; Histograms of the number of place fields per neuron and distributions of field sizes. Right; Distributions of field sizes. Individual fields are shown as open circles and means are shown as lines. Generalized linear mixed effects model testing the effects of experimental group on number of fields for each region separately with mouse and neuron nested in mouse as random factors. CA1: No effect of group on number of fields per cell (control vs. KA t(122) = -0.70, p = 0.487; estimated means: control = 2.26 +/- 0.25, KA = 1.99, +/- 0.29). CA3: No effect of group on number of fields per cell (control vs. KA t(83) = 0.29, p = 0.770; estimated means: control = 1.90 +/- 0.26, KA = 1.80, +/- 0.23). DG: No effect of group on number of fields per cell (control vs. KA t(35) = -1.23, p = 0.229; estimated means: control = 1.86 +/- 0.34, KA = 2.39, +/- 0.27). Generalized linear mixed effects model testing the effects of experimental group on field size for each region separately with mouse and field nested in mouse as random factors (field treated independently). CA1: No effect of group on field size (control vs KA t(265) = 0.07, p = 0.941; estimated means: control = 256 +/- 42.1, KA = 251 +/- 48.1). CA3: No effect of group on field size (control vs KA t(152) = 1.26, p = 0.210; estimated means: control = 246 +/- 41.3, KA = 176 +/- 37.0). DG: No effect of group on field size (control vs KA t(79) = 0.59, p = 0.558; estimated means: control = 223 +/- 55.2, KA = 184 +/- 35.8).

**Figure S3. Cross region coherence in control and epileptic mice.** Coherence was calculated in three oscillatory bands (2-4 Hz, 6-12 Hz, and 14-20 Hz) between CA1 and CA3,

CA1 and DG, and DG and CA3. Values from individual mice are shown as circles. Means are shown as bars. Generalized linear mixed effects model testing the effects of experimental group on coherence for each region pair, with p-values uncorrected and corrected for multiple comparisons (3 per region pair). Nothing is significant after correction, however trends toward higher coherence in KA mice for CA1/CA3 were observed in two frequency bands.

| Region pair | Frequency band | df | t-value | Uncorrected p-value | Corrected p-value |
|---|---|---|---|---|---|
| CA1/CA3 | 2-4 Hz | 12 | -1.66 | 0.124 | 0.372 |
| CA1/CA3 | 6-12 Hz | 12 | -2.12 | 0.0559 | 0.167 |
| CA1/CA3 | 14-20 Hz | 12 | -2.71 | 0.0191 | 0.0573 |
| CA1/DG | 2-4 Hz | 9 | 0.31 | 0.763 | 1 |
| CA1/DG | 6-12 Hz | 9 | 1.64 | 0.136 | 0.408 |
| CA1/DG | 14-20 Hz | 9 | 1.35 | 0.210 | 0.630 |
| CA3/DG | 2-4 Hz | 9 | 0.86 | 0.414 | 1 |
| CA3/DG | 6-12 Hz | 9 | 0.46 | 0.654 | 1 |
| CA3/DG | 14-20 Hz | 9 | 0.34 | 0.741 | 1 |

**Figure S4. Interictal spike rates and measures of spatial coding in epileptic mice. A,** A representative trace from CA1 during foraging showing frequent interictal spikes. **B,** The average spatial information, spatial coherence, and spatial correlations of place cells for CA1 (top) and CA3 (bottom) plotted as a function of average interictal spike rate. Each dot is an animal. Only one relationship was significantly correlated; IS rate vs. CA1 spatial information, $p = 0.010$.

**Figure S5. Pathohistology and measures of spatial coding in epileptic mice. A,** Examples of histological sections with measures of granule cell layer width and CA3 pyramidal cell layer area shown for every epileptic animal included in the study. Note that for each animal, 3 sections were analyzed (the averages of the measures are shown in Table 1). **B,** DG dispersion and CA3 area were significantly correlated ($p = 0.05$). For additional testing, we used the measure of DG dispersion as we felt more confident in that measure. **C,** There was no correlation observed between DG dispersion and average spatial information, spatial coherence, and spatial correlation for CA1 (top) or CA3 (bottom).

**Statistical Companion**

**Figure 1A,** Generalized linear mixed effects model testing the effect of experimental group on peak firing rate. Animal and cell nested within animal were included as random factors. No significant effect of experimental group for CA1 (control vs. KA, t(175) = 0.04, p = 0.969; estimated means: control (n = 111) = 12.1 +/- 2.00, KA (n = 66) = 12.0 +/- 1.92), CA3 (control vs. KA, t(155) = 1.92, p = 0.0565; estimated means: control (n = 61) = 13.9 +/- 1.34, KA (N = 96) = 10.5 +/- 1.07), or DG (control vs. KA, t(75) = 0.47, p = 0.643; estimated means: control (n = 26) = 10.0 +/- 3.17, KA (n = 51) = 8.2 +/- 2.06). **B,** No significant effect of experimental group for CA1 (control vs. KA, t(175) = 0.750, p = 0.454; estimated means: control = 0.61 +/- 0.118, KA = 0.48 +/- 0.119), CA3 (control vs. KA, t(155) = -0.753, p = 0.453; estimated means: control = 0.61 +/- 0.128, KA = 0.73 +/- 0.102), or DG (control vs. KA, t(75) = 0.160, p = 0.874; estimated means: control = 0.45 +/- 0.108, KA = 0.42 +/- 0.167). The proportion of place cells in CA1 of KA mice was significantly smaller than in controls ($\chi^2$ = 12.01, $p \leq 0.001$). Significance denoted with asterisks: ***p ≤ 0.001. There was no significant difference in the proportion of place cells between control and KA mice for CA3 ($\chi^2$ = 0.20, $p$ = 0.66) and DG ($\chi^2$ = 0.13, $p$ = 0.72). Wilcoxon rank sum test and Chi-squared test.

**Figure 2B,** Generalized linear mixed effects model testing the effect of experimental group on half map spatial correlation for place cells. Animal and cell nested within animal were included as random factors. No significant effect of experimental group for CA1 (control vs. KA t(122) = 0.07, p = 0.942; estimated means: control (n = 86) = 0.82 +/- 0.0199, KA (n = 38) = 0.820 +/- 0.268*)*. Significant effect of experimental group for CA3 (control vs. KA t(83) = 2.1, p = 0.0350; estimated means: control (n = 37) = 0.88 +/- 0.032; KA (n = 48) = 0.79 +/- 0.0281). No significant effect of experimental group for DG (control vs. KA t(35) = 1.43, p = 0.161; estimated means: control (n = 14) = 0.87 +/- 0.0533, KA (n = 23) = 0.78 +/- 0.0396). **C,** Generalized linear mixed effects model testing the effect of experimental group on coherence for place cells. Significant effect of experimental group for CA1 (control vs. KA t(122) = 2.24, p = 0.027; estimated means: control (n = 86) = 0.55 +/- 0.040, KA (n = 38) = 0.42 +/- 0.044). No significant effect of experimental group for CA3 (control vs. KA t(83) = 1.5, p = 0.137; estimated means: control (n = 37) = 0.56 +/- 0.0552, KA (n = 48) = 0.46 +/- 0.0479. No significant effect of experimental group for DG (control vs. KA t(35) = 1.2, p = 0.247; estimated means: control (n = 14) = 0.55 +/- 0.0423, KA (n = 23) = 0.48 +/- 0.0353).

Other features:

CA1:

| *Feature* | *d.f.* | *t-value* | *p-value* | *Control est. mean* | *Control SEM* | *KA est. mean* | *KA SEM* |
|---|---|---|---|---|---|---|---|
| peak rate | 122 | -0.137 | 0.891 | 12.7 | 2.03 | 13.1 | 2.21 |
| average rate | 122 | 0.493 | 0.623 | 2.9 | 0.28 | 2.7 | 0.37 |
| information | 122 | -0.174 | 0.862 | 0.74 | 0.118 | 0.77 | 0.142 |
| sparsity | 122 | 0.590 | 0.557 | 0.53 | 0.050 | 0.49 | 0.056 |
| selectivity | 122 | -0.657 | 0.512 | 7.4 | 1.47 | 8.9 | 1.87 |

CA3:

| *Feature* | *d.f.* | *t-value* | *p-value* | *Control est. mean* | *Control SEM* | *KA est. mean* | *KA SEM* |
|---|---|---|---|---|---|---|---|
| peak rate | 83 | 0.768 | 0.444 | 15.3 | 1.83 | 13.4 | 1.60 |
| average rate | 83 | 1.23 | 0.222 | 3.0 | 0.36 | 2.4 | 0.31 |
| information | 83 | -1.27 | 0.209 | 0.89 | 0.188 | 1.21 | 0.162 |
| sparsity | 83 | 1.03 | 0.308 | 0.48 | 0.058 | 0.40 | 0.050 |
| selectivity | 83 | -1.00 | 0.317 | 10.2 | 2.44 | 13.5 | 2.11 |

DG:

| *Feature* | *d.f.* | *t-value* | *p-value* | *Control est. mean* | *Control SEM* | *KA est. mean* | *KA SEM* |
|---|---|---|---|---|---|---|---|
| peak rate | 35 | 0.501 | 0.619 | 14.9 | 2.77 | 13.2 | 2.16 |
| average rate | 35 | -0.458 | 0.650 | 2.6 | 0.553 | 2.9 | 0.432 |
| information | 35 | 1.41 | 0.169 | 0.92 | 0.151 | 0.65 | 0.118 |
| sparsity | 35 | -0.830 | 0.412 | 0.45 | 0.059 | 0.51 | 0.046 |
| selectivity | 35 | 2.29 | 0.0283 | 11.0 | 1.61 | 6.33 | 1.26 |

**Figure 3C,**

CA1: Generalized linear mixed effects models testing the effect of comparison combination on spatial correlation for each group separately (i.e., comparing F1/F2 vs F1/N1 for control mice and KA mice separately). Significant effect of combination for control mice (t(140) = 6.93, p = 1.43 e-10; estimated means: f1/f2 (n = 71) = 0.58 /- 0.040, f1/n1 (n = 71) = 0.21 +/- 0.040). Significant effect of combination for KA mice (t(48) = 6.62, p = 2.848 e-8; estimated means: f1/f2 (n = 24) = 0.64 +/- 0.082, f1/n1 (n = 27) = 0.09 +/- 0.080). Generalized linear mixed effects model testing the effect of group on F1/F2 stability on spatial correlation. No significant effect of group (control vs KA) (t(93) = 0.68, p = 0.499; estimated means: control (n = 71) = 0.58 +/- 0.038, KA (n = 24) = 0.63 +/- 0.066).

CA3: Generalized linear mixed effects models testing the effect of comparison combination on spatial correlation for each group separately (i.e., comparing F1/F2 vs F1/N1 for control mice and KA mice separately). Significant effect of combination for control mice (t(64) = 5.42, p = 9.82 e -7; estimated means: f1/f2 (n = 33) = 0.76 +/- 0.084, f1/n1 (n = 33) = 0.34 +/- 0.083). Significant effect of combination for KA mice (t(82) = 8.24, p = 2.36 e-12; estimated means: f1/f2 (n = 44) = 0.63 +/- 0.073, f1/n1 (n = 42) = 0.09 +/- 0.0073). Generalized linear mixed effects model testing the effect of group on F1/F2 stability on spatial correlation. No significant effect of group (control vs KA, (t(74) = 1.71, p = 0.0906; estimated means: control (n = 33) = 0.81 +/- 0.114, KA (n = 44) = 0.57 +/- 0.085).

DG: Generalized linear mixed effects models testing the effect of comparison combination on spatial correlation for each group separately (i.e., comparing F1/F2 vs F1/N1 for control mice and KA mice separately). Significant effect of combination for control mice (t(20) = 5.55, p = 1.96 e-5; estimated means: f1/f2 (n = 11) = 0.80 +/- 0.102, f1/n1 (n = 11) = 0.002 +/- 0.102). Significant effect of combination for KA mice (t(51) = 11.18, p = 2.53 e-15; estimated means: f1/f2 (n = 27) = 0.79 +/- 0.066, f1/n1 (n = 26) = 0.08 +/- 0.067). Generalized linear mixed effects model testing the effect of group on F1/F2 stability on spatial correlation. No significant effect of group (control vs KA) (t(36) = 0.78, p = 0.441; estimated means: control (n = 11) = 0.80 +/- 0.055, KA (n = 27) = 0.75 +/- 0.035).

**E,** CA1: Generalized linear mixed effects model testing the effects of experimental group on N1/N4 spatial correlation scores. No significant effect of group (t(94) = -0.78, p = 0.440; estimated means: control (n = 69) = 0.43 +/- 0.059, KA (n = 27) = 0.50 +/- 0.077).

CA3: Generalized linear mixed effects model testing the effects of experimental group on N1/N4 spatial correlation scores. No significant effect of group (t(64) = -1.90, p = 0.062; estimated means: control (n = 27) = 0.67 +/- 0.113, KA (n = 39) = 0.41 +/- 0.083).

DG: Generalized linear mixed effects model testing the effects of experimental group on N1/N4 spatial correlation scores. No significant effect of group (t(34) = 0.45, p = 0.655; estimated means: control (n = 11) = 0.49 +/- 0.209, KA (n = 25) = 0.38 +/- 0.132).

**Figure S1**:

Generalized linear mixed effects model testing the effect of experimental group on max average firing rate for principal cells, tested for each region separately. No significant effect of experimental group for CA1 principal cell firing rate (control vs. KA, t(175) = -1.0, p = 0.319; estimated means: control (n = 111) = 3.38 +/- 0.585, KA (n = 66) = 4.20 +/- 0.561). Significant effect of experimental group for CA3 (control vs. KA, t(155) = 2.01, p = 0.0457; estimated means: control (n = 61) = 4.02 +/- 0.348, KA (n = 96) = 3.12 +/- 0.277. No significant effect of experimental group for DG (control vs. KA t(75) = -0.446, p = 0.657; estimated means: control (n = 26) = 3.09 +/- 0.481, KA (n = 51) = 2.70 +/- 0.740.

Generalized linear mixed effects model testing the effect of experimental group on max average firing rate for interneurons, tested for each region separately. No significant effect of experimental group for CA1 interneuron firing rate (control vs. KA, t(50) = -2.0, p = 0.0541; estimated means: control (n = 32) = 20.14 +/- 3.61, KA (n = 20) = 30.64 +/- 3.91). No significant effect of experimental group for CA3 interneuron firing rate (control vs. KA, t(62) = 0.65, p = 0.520; estimated means: control (n = 23) = 27.1 +/- 2.78), KA (n = 41) = 24.84 +/- 2.08). No significant effect of experimental group for DG interneuron firing rate (control vs. KA, t(18) = 0.809, p = 0.429; estimated means: control (n = 6) = 34.3 +/- 6.74, KA (n = 14) = 23.6 +/- 11.4).

Generalized linear mixed effects model testing the effect of experimental group on burst index for principal cells, tested for each region separately. No significant effect of experimental group for CA1 principal cell firing rate (control vs. KA t(173) = -0.522, p = 0.603; estimated means:

control (n = 111) = 0.35 +/- 0.053, KA (n = 66) = 0.39 +/- 0.048). No significant effect of experimental group for CA3 principal cell firing rate (control vs. KA t(151) = -1.84, p = 0.0683; estimated means: control (n = 61) = 0.32 +/- 0.050, KA (n = 96) = 0.44 +/- 0.040). No significant effect of experimental group for DG principal cell firing rate (control vs. KA t(74) = 0.55, p = 0.585; estimated means: control (n = 26) = 0.32 +/- 0.050, KA (n = 51) = 0.44 +/- 0.040).

**Figure S2:**

**B,** Generalized linear mixed effects model testing the effects of experimental group on number of fields for each region separately with mouse and unit nested in mouse as random factors.

CA1: No effect of group on number of fields per cell (control vs. KA t(122) = -0.70, p = 0.487; estimated means: control (n = 86) = 2.26 +/- 0.25, KA (n = 38) = 1.99, +/- 0.29).

CA3: No effect of group on number of fields per cell (control vs. KA t(83) = 0.29, p = 0.770; estimated means: control (n = 37) = 1.90 +/- 0.26, KA (n = 48) = 1.80, +/- 0.23).

DG: No effect of group on number of fields per cell (control vs. KA t(35) = -1.23, p = 0.229; estimated means: control (n = 14) = 1.86 +/- 0.34, KA (n = 23) = 2.39, +/- 0.27).

**C,** Generalized linear mixed effects model testing the effects of experimental group on field size for each region separately with mouse and field nested in mouse as random factors (field treated independently).

CA1: No effect of group on field size (control vs KA t(265) = 0.07, p = 0.941; estimated means: control (n = 194) = 256 +/- 42.1, KA (n = 73) = 251 +/- 48.1).

CA3: No effect of group on field size (control vs KA t(152) = 1.26, p = 0.210; estimated means: control (n = 69) = 246 +/- 41.3, KA (n = 85) = 176 +/- 37.0).

DG: No effect of group on field size (control vs KA t(79) = 0.59, p = 0.558; estimated means: control (n = 26) = 223 +/- 55.2, KA (n = 55) = 184 +/- 35.8).

**Figure S3:**

Generalized linear mixed effects model testing the effects of experimental group on coherence for each region pair, with p-values uncorrected and corrected for multiple comparisons (3 per region pair). Nothing is significant after correction.

| *Region pair* | *Frequency band* | *df* | *t-value* | *Uncorrected p-value* | *Corrected p-value* |
|---|---|---|---|---|---|
| CA1/CA3 | 2-4 Hz | 12 | -1.66 | 0.124 | 0.372 |
| CA1/CA3 | 6-12 Hz | 12 | -2.12 | 0.0559 | 0.167 |
| CA1/CA3 | 14-20 Hz | 12 | -2.71 | 0.0191 | 0.0573 |
| CA1/DG | 2-4 Hz | 9 | 0.31 | 0.763 | 1 |

| | | | | | |
|---|---|---|---|---|---|
| CA1/DG | 6-12 Hz | 9 | 1.64 | 0.136 | 0.408 |
| CA1/DG | 14-20 Hz | 9 | 1.35 | 0.210 | 0.630 |
| CA3/DG | 2-4 Hz | 9 | 0.86 | 0.414 | 1 |
| CA3/DG | 6-12 Hz | 9 | 0.46 | 0.654 | 1 |
| CA3/DG | 14-20 Hz | 9 | 0.34 | 0.741 | 1 |

| *Region pair* | *Frequency band* | *Control est. mean* | *Control SEM* | *KA est. mean* | *KA SEM* |
|---|---|---|---|---|---|
| CA1/CA3 | 2-4 Hz | 0.60 | 0.042 | 0.69 | 0.037 |
| CA1/CA3 | 6-12 Hz | 0.70 | 0.031 | 0.78 | 0.027 |
| CA1/CA3 | 14-20 Hz | 0.57 | 0.039 | 0.70 | 0.034 |
| CA1/DG | 2-4 Hz | 0.60 | 0.075 | 0.57 | 0.046 |
| CA1/DG | 6-12 Hz | 0.79 | 0.068 | 0.66 | 0.042 |
| CA1/DG | 14-20 Hz | 0.67 | 0.047 | 0.60 | 0.029 |
| CA3/DG | 2-4 Hz | 0.77 | 0.108 | 0.66 | 0.066 |
| CA3/DG | 6-12 Hz | 0.77 | 0.086 | 0.72 | 0.053 |
| CA3/DG | 14-20 Hz | 0.70 | 0.096 | 0.66 | 0.059 |

**Figures S4** and **S5** show statistics performed on individual animals rather than cells and p values reported the figures.
